# RECON gene disruption enhances host resistance to enable genome-wide evaluation of intracellular pathogen fitness during infection

**DOI:** 10.1101/2024.01.15.575726

**Authors:** Chelsea E. Stamm, Adelle P. McFarland, Melissa N. Locke, Hannah Tabakh, Qing Tang, Maureen K. Thomason, Joshua J. Woodward

**Affiliations:** Department of Microbiology, University of Washington, Seattle, WA, USA; Molecular and Cellular Biology Program, University of Washington, Seattle, WA, USA

## Abstract

Transposon sequencing (Tn-seq) is a powerful genome-wide technique to assess bacterial fitness under varying growth conditions. However, screening via Tn-seq *in vivo* is challenging. Dose limitations and host restrictions create bottlenecks that diminish the transposon mutant pool being screened. Here we have developed a murine model with a disruption in *Akr1c13* that renders the resulting RECON^-/-^ mouse resistant to high dose infection. We leveraged this model to perform a Tn-seq screen of the human pathogen *Listeria monocytogenes in vivo*. We identified 139 genes which were required for *L. monocytogenes* growth in mice including novel genes not previously identified for host survival. We identified organ specific requirements for *L. monocytogenes* survival and investigated the role of the folate enzyme FolD in *L. monocytogenes* liver pathogenesis. A mutant lacking *folD* was impaired for growth in murine livers by 2.5-log_10_ compared to wildtype and failed to spread cell-to-cell in fibroblasts. In contrast, a mutant in *alsR,* which encodes a transcription factor that represses an operon involved in D-allose catabolism, was attenuated in both livers and spleens of mice by 4-log_10_ and 3-log_10_, respectively, but showed modest phenotypes in *in vitro* models. We confirmed that dysregulation of the D-allose catabolism operon is responsible for the *in vivo* growth defect, as deletion of the operon in the Δ*alsR* background rescued virulence. By undertaking an unbiased, genome-wide screen in mice, we have identified novel fitness determinants for *L. monocytogenes* host infection, which highlights the utility of the RECON^-/-^ mouse model for future screening efforts.

**Importance:** *Listeria monocytogenes* is the gram-positive bacterium responsible for the food-borne disease Listeriosis. Although infections with *L. monocytogenes* are limiting in healthy hosts, vulnerable populations, including pregnant and elderly people, can experience high rates of mortality. Thus, understanding the breadth of genetic requirements for *L. monocytogenes in vivo* survival will present new opportunities for treatment and prevention of Listeriosis. We developed a murine model of infection using a RECON^-/-^ mouse that is restrictive to systemic *L. monocytogenes* infection. We utilized this model to screen for *L. monocytogenes* genes required *in vivo* via Tn-seq. We identified the liver-specific gene *folD* and a repressor, *alsR* that only exhibits an *in vivo* growth defect. AlsR controls the expression of the D-allose operon which is a marker in diagnostic techniques to identify pathogenic Listeria. A better understanding of the role of the D-allose operon in human disease may further inform diagnostic and prevention measures.

## Introduction

*Listeria monocytogenes* (Lm) is a gram-positive, facultative pathogen and is the main agent responsible for the food-borne disease Listeriosis. Lm is ubiquitous in the environment and can grow under conditions used in food processing such as high salt and low temperatures (1). Asymptomatic ingestion of contaminated food by immune competent individuals is predicted to occur several times a year (2). However, exposure among vulnerable populations such as pregnant people, the immunocompromised and the elderly frequently causes invasive disease with severe morbidity and mortality (3, 4). In these individuals, the liver and spleen are replicative niches where Lm grows to high numbers and can further disseminate across the blood-brain or placental barriers to cause severe disease (5–7). Thus, understanding the genetic requirements for Lm to replicate in the liver and spleen may provide novel treatment targets for prevention of severe disease.

Lm pathogenesis is facilitated by its ability to infect multiple cell types intracellularly. The Lm intracellular life cycle is well-established and relies on virulence genes clustered together on the Listeria Pathogenicity Island-I (LPI), which are under the control of PrfA (4). Functions for endosomal escape, actin-based motility, and cell-to-cell spread are all encoded by genes on the LPI (8). In addition to virulence gene expression, intracellular replication requires metabolic reprogramming to withstand nutrient restriction in the cytosol (9, 10). While much of the intracellular life cycle of Lm and virulence properties of the LPI were comprehensively determined using cell culture methods, discrepancies exist among *in vitro* models. For example, the actin nucleating protein ActA is required for development of plaques in epithelial cells, but it is dispensable for intracellular replication in macrophage infections (11). In addition, binding of the internalin InlB to non-phagocytic cells in cell culture is promiscuous and in opposition to observed tropism *in vivo* where it is not required for Lm intestinal barrier crossing but for dissemination to peripheral organs (12, 13). Finally, it is clear that homogenous cell lines cannot replicate the complex host-pathogen interactions inherent in a whole organism (14). Therefore, establishing the genetic requirements of Lm survival *in vivo* is of utmost interest.

Transposon mutagenesis has long facilitated genotype-to-phenotype discoveries in bacteria but until recently was hampered by low-throughput screening methods. Transposon sequencing (Tn-seq) leverages next-generation sequencing to assess the fitness of a pool of transposon-mutagenized bacteria under a given selection in a high-throughput manner (15, 16). Tn-seq technologies have greatly increased the genetic landscape that can be assessed in a single experiment and have been used to establish essential genes and genes required for host survival for several pathogenic bacteria of interest including *Streptococcus pneumoniae* (15), *Vibrio cholerae* (17), *Acinetobacter baumannii* (18) and *Staphylococcus aureus* (19, 20). However, the technical challenges associated with utilizing Tn-seq *in vivo* including bottlenecks of the transposon pool that arise from host restrictions and the potential cost prohibitive nature of screening in animals limits Tn-seq broader usage (16). For example, the low founding population in mouse peripheral organs at a lethal dose 50 (LD50) of Lm restricts the total number of transposon mutants that can be delivered to each animal (21). Here, we describe the development of a mouse model lacking a single protein, RECON^-/-^, that can survive increased bacterial burdens. Using Lm as a model pathogen, we show that this mouse is a viable model for assessing bacterial fitness factors *in vivo* via Tn-seq. Finally, we validate two hits, *folD* and *alsR* in wildtype mice and characterize their roles in Lm *in vivo* pathogenesis.

## Results

### Generation of RECON-deficient mice

We previously identified a murine aldo-keto reductase (AKR) encoded by *Akr1c13* that binds the bacterial second messenger cyclic diadenosine monophosphate (c-di-AMP). Binding of the protein RECON to c-di-AMP inhibits its enzymatic activity and results in augmented NF-κB activation (22). To better understand the role of RECON in the immune response to bacterial infection, we used CRISPR/Cas9 to mutagenize *Akr1c13* in mice. *In vitro* transcribed sgRNA and Cas9 mRNA were microinjected into C57BL/6J embryos and implanted into pseudo-pregnant female mice (Figure S1A). We selected a heterozygous male founder that had a single G insertion in exon 6 of *Akr1c13* (Figures S1A and S1B) and bred the line to homozygosity. The single nucleotide insertion in the *Akr1c13* coding sequencing caused a frameshift resulting in a premature stop codon after residue 200, which truncates the protein prior to the active site and c-di-AMP binding site (Figures 1A and S1B). Due to close homology with other murine AKRs, there is not a RECON-specific antibody. However, we were able to analyze the expression of *Akr1c13* in liver, spleen, lung, and bone marrow derived macrophages (BMDM) via quantitative RT-PCR (qRT-PCR). We observed that RNA levels of *Akr1c13* decreased in the CRISPR/Cas9-targeted mice compared to wild-type mice (Figure 1B), likely due to nonsense-mediated decay of unstable mRNAs (23). Therefore, we concluded these mice are deficient in RECON (RECON^-/-^) as a result of a frameshift mutation that reduces mRNA stability and abolishes expression of the full-length protein.

**Figure 1.**
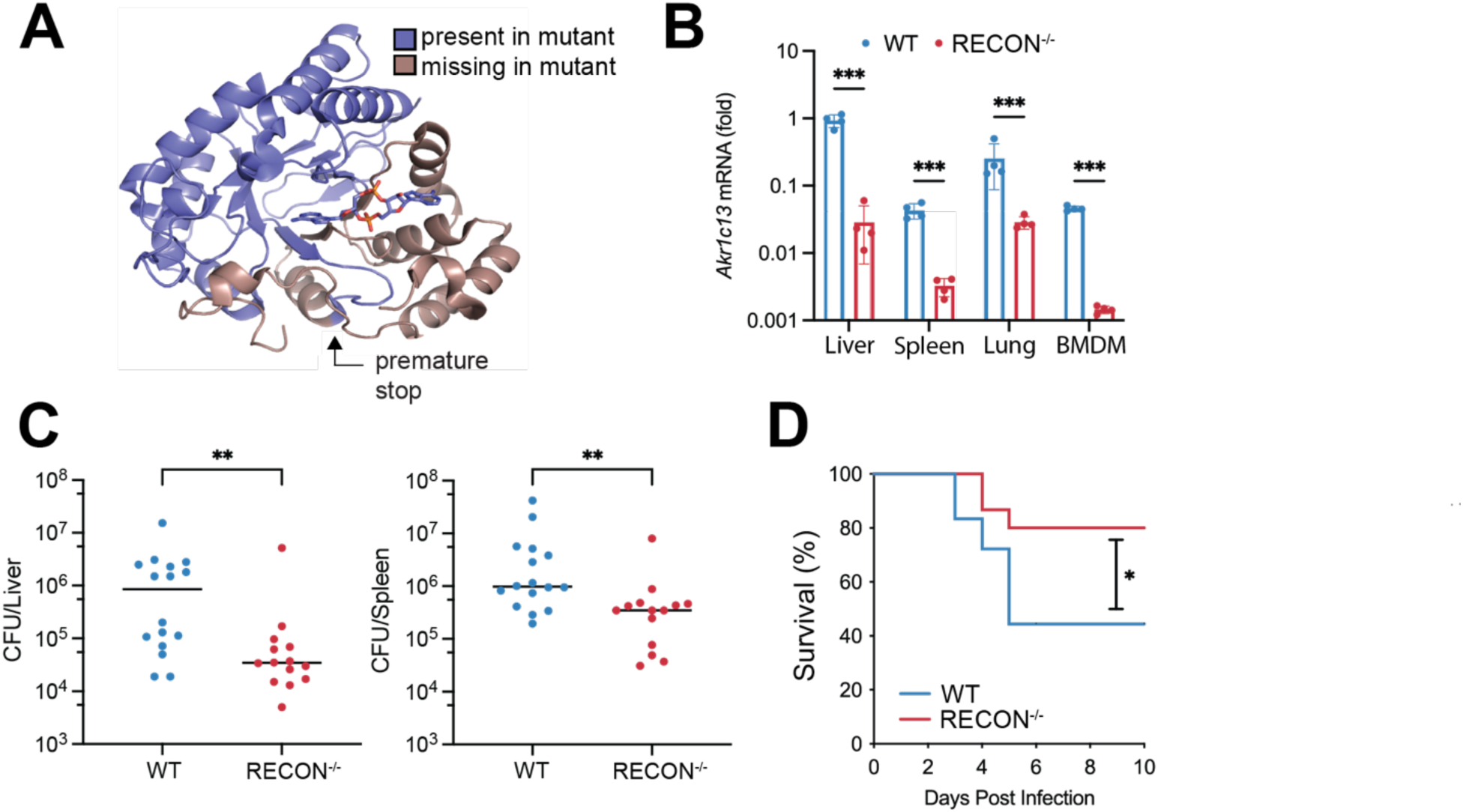
Loss of RECON leads to bacterial clearance. (**A**) RECON protein structure with the region missing in the truncated mutant protein colored in brown. Truncation destroys the active and c-di-AMP binding sites. (**B**) qRT-PCR analysis on mRNA from mouse liver, spleen, lung, and BMDM isolated from WT and RECON^-/-^ mice (N=4 for each genotype). The endogenous control gene was *Hprt* and data are normalized to the mean value of *Akr1c13*/*Hprt* from livers of WT mice. Median values are indicated by a bar. Statistical significance was determined by an unpaired Student’s t test with ***p <0.001. (**C**) WT or RECON*^-/-^* mice were injected IP with 1×10^4^ CFU Lm. CFU were enumerated from liver (left) and spleen (right) 72 hours post infection. Data are combined from two and are representative of more than three independent experiments. (**D**) Kaplan-Meier survival curve shown for WT (n=18) or RECON^-/-^ (n=15) mice injected IP with 1×10^6^ CFU Lm. For CFU enumeration (**C**) and survival (**D**), statistical significance was assessed by Mann-Whitney U test and log-rank test, respectively. *p<0.05, **p<0.005.

### RECON-deficient mice are more resistant to bacterial infection

As we had previously demonstrated that altering expression levels of RECON affected the intracellular survival of Lm in macrophages but not hepatocytes (22), we sought to determine how loss of RECON would influence the outcome of Lm infection *in vivo*. To that end, we infected WT or RECON-deficient littermates with Lm by intraperitoneal (IP) injection and determined bacterial burden at 72 hours post infection (hpi). Bacterial colony forming units (CFU) collected from livers and spleens of RECON^-/-^ mice were significantly lower than in wild-type littermates (Figure 1C). To determine if this decrease in CFU resulted in protection for RECON^-/-^ mice, we again infected Lm intraperitoneally and monitored murine survival over the course of ten days. Consistent with the decreased bacterial burden, RECON^-/-^ mice exhibited increased survival to Lm infection compared to wild-type mice (Figure 1D). Together these data demonstrate that loss of RECON promotes faster bacterial clearance during systemic challenge and that bacterial restriction is protective.

### RECON-deficient mice are a viable model for *in vivo* screening by transposon sequencing

As previously discussed, technical limitations impede the use of Tn-seq *in vivo*. However, since RECON^-/-^ mice were more resistant to Lm infection (Figure 1D-E), we hypothesized that this mouse genotype would survive acute infection with a high dose inoculum, circumventing some of the technical challenges associated with Lm Tn-seq and be a powerful animal model to assess bacterial fitness using Tn-seq *in vivo*.

To that end, we generated a *mariner*-based transposon library in Lm strain 10403S containing 17,000 mutants and infected RECON^-/-^ mice with 2.4 x 10^6^ CFU/mouse via an intravenous (IV) route. We collected bacteria from livers and spleens 34 hpi, isolated genomic DNA and prepared libraries for massive parallel sequencing of the transposon insertion junctions. Then, we used PATRIC (24) to identify genes that were significantly (p<0.05) depleted in livers or spleens of mice compared to the input transposon library (Figures 2A and 2B, respectively). We identified 124 genes in the liver and 45 genes in the spleen with at least a two-fold decrease in reads compared to the input (Tables S1 and S2, respectively). Importantly, reads in *prfA*, *hly*, *actA* and *plcB* which are components of the LPI, and required for Lm virulence (4), were absent in all but two liver samples (Figure 2C). In fact, the LPI genes were among the most significantly depleted genes in both organs (Figures 2A and 2B). In total, we identified 34 genes that were required for growth in both organs (Figure 2D), many of which have been previously implicated in virulence (Tables S1 and S2) which confirms the robustness of our screening approach.

**Figure 2.**
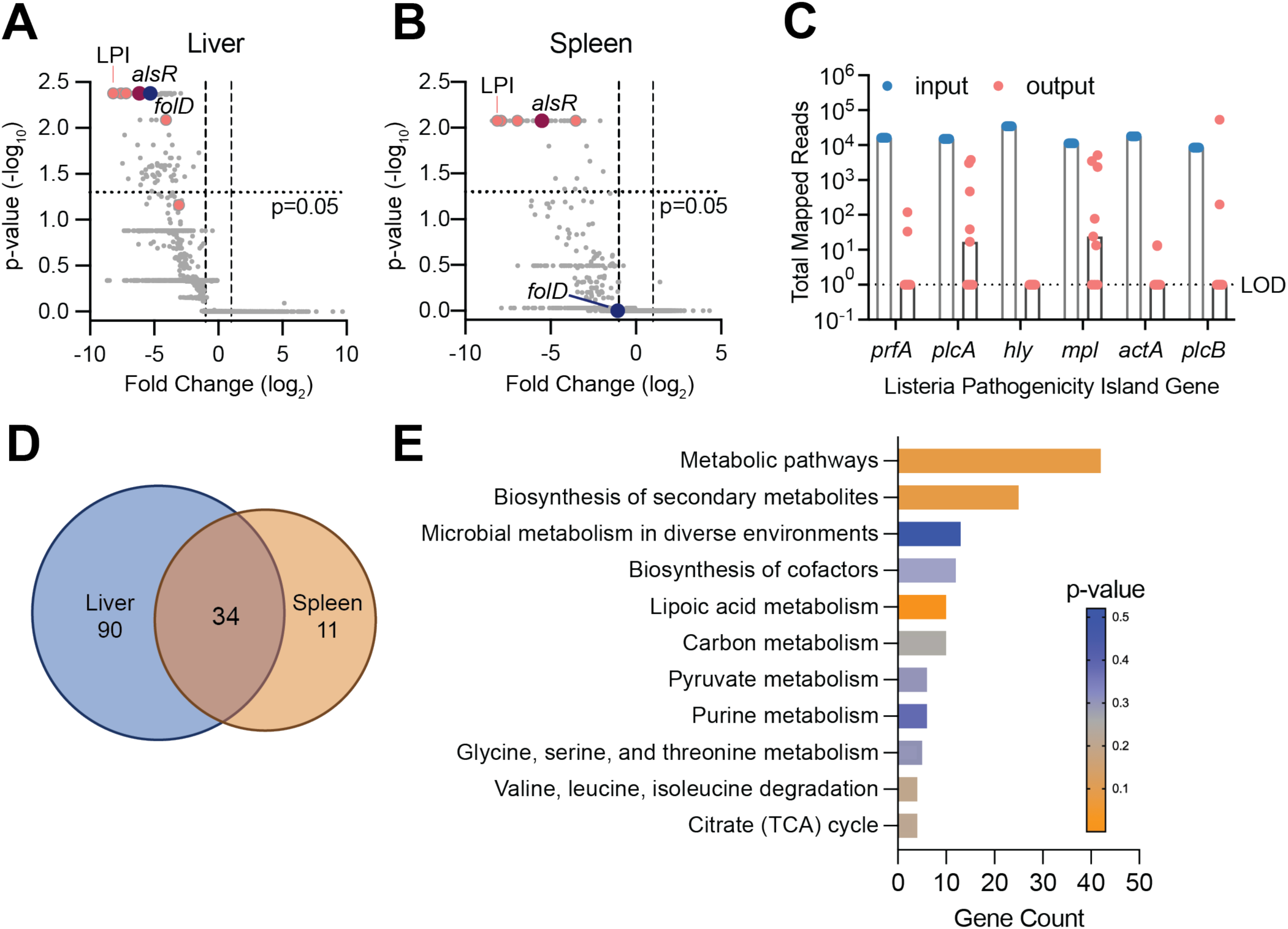
Identification of Lm genes required for murine infection via TN-Seq. (**A-B**) Volcano plots of mapped Lm genes from livers (**A**) and spleens (**B**) of infected RECON^-/-^ mice. Each dot represents a single gene and vertical lines denote two-fold change. The LPI is highlighted in pink and two hits, *alsR* and *folD,* are indicated by red and blue, respectively. (**C**) Total mapped reads for each gene in the LPI in the input library compared to the liver output library. Data for each mouse is plotted as a dot and the median is indicated by the bars. LOD, limit of detection. (**D**) Venn diagram comparing the 124 genes required for Lm infection of the liver with the 45 genes required in the spleen. (**E**) KEGG pathway enrichment analysis of all hits.

We analyzed the genes required for growth *in vivo* by pathway enrichment analysis (Figure 2E). Lm is an auxotroph for lipoic acid and genes required for lipoic acid acquisition were the most significantly enriched. However, the majority of enriched genes clustered into the general metabolic pathways group and could be further broken down into genes enriched for energy production (carbon metabolism, pyruvate metabolism and citrate (TCA) cycle). In addition, genes required for *in vivo* survival were enriched in pathways for the synthesis of secondary metabolites, essential cofactors, purines, and amino acids. Finally, nearly 50% of the genes required for *in vivo* survival were not significantly enriched in KEGG pathways including a proportion of the genes (9%) that remained unannotated and could not be mapped to any pathway. Taken together, these data confirm that core metabolic processes are indispensable for growth in the murine host and suggests that there are genes with *in vivo*-specific functions yet to be elucidated.

### The folate metabolism gene *folD* contributes to *L. monocytogenes* fitness in the murine liver

The primary niches for Lm in the murine liver and spleen are distinct, with hepatocytes being the main cell type for Lm intracellular growth in the liver (25, 26). To better understand the genetic requirements for liver colonization, we further investigated the unique genes required for Lm growth in the liver only. One such gene, *folD* (*lmo1360*) which encodes the bifunctional enzyme 5,10-methylene-tetrahydrofolate dehydrogenase/ 5,10-methylene-tetrahydrofolate cyclohydrolase, was depleted nearly 40 fold in livers of mice but was not significantly changed in the spleens (Figures 2A and 2B and Table S1).

The bioactive tetrahydrofolates (THF) produced by FolD in the one-carbon cycle are used to synthesize deoxythymidine monophosphate (dTMP), purines and the initiator amino acid N-formylmethionine (Figure 3A) (27, 28). To validate the organ-specificity of *folD*, we deleted it from the Lm 10403S chromosome, infected WT C57BL/6 mice via IV and harvested CFU from livers and spleens at 72 hpi. We recovered over two logs fewer Δ*folD* bacteria than WT from livers of mice, which could be rescued by constitutive expression of *folD* from a separate chromosomal locus (Δ*folD*::*folD*) (Figure 3B). In contrast, Δ*folD* growth in the spleen was only reduced by half a log compared to WT and was similarly rescued by complementation (Figure 3C). We confirmed the virulence defect of Δ*folD* in the murine liver was not due to an overall replication defect by measuring growth over time in brain heart infusion (BHI) and Listeria defined (29) minimal media (MM) and observed no difference in growth between WT and Δ*folD* in either condition (Figure S2A and S2B).

**Figure 3.**
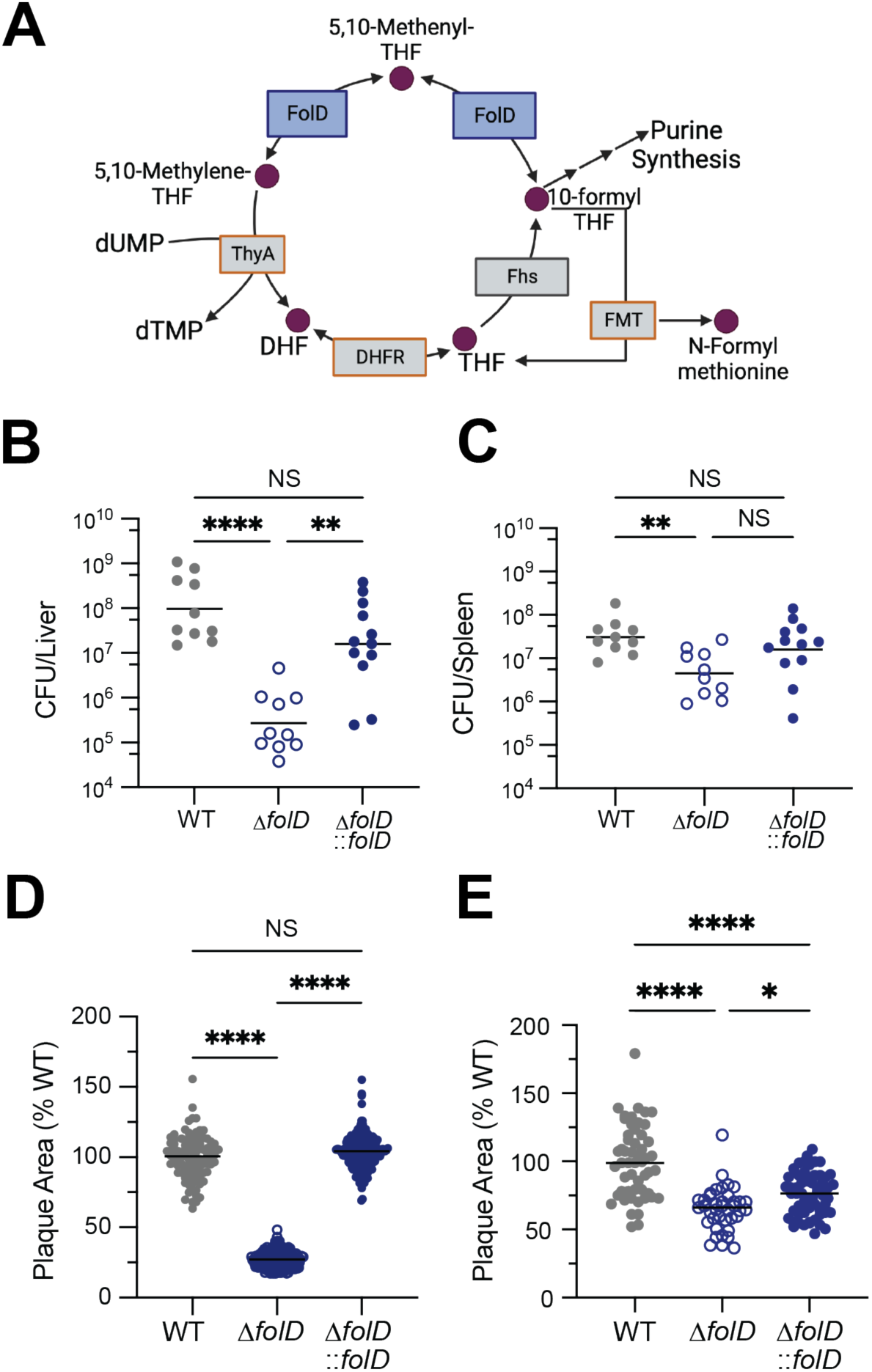
The folate metabolism gene *folD* contributes to Lm fitness in the murine liver. (**A**) Schematic of folate cycle in Lm. FolD is indicated in blue. Genes outlined in orange were essential in our library. (**B-C**) CFU harvested from livers (**B**) and spleens (**C**) of WT mice infected with WT, Δ*folD* or Δ*folD*::*folD* Lm for 72 hours. Data are combined from two independent experiments. ** p<0.01, **** p<0.0001 by Kruskal-Wallis test. (**D-E**) Plaque area measured in fibroblasts (**D**) or hepatocytes (**E**) that were infected with WT, Δ*folD* or Δ*folD*::*folD* Lm and stained with neutral red after 48 h. Data are combined from triplicate wells and are representative of two independent experiments. * p<0.05, **** p< 0.0001 by nonparametric ANOVA.

Lm replicates to high CFU in the liver in part due to its ability to spread cell-to-cell, without encountering the extracellular space (30, 31). In order to determine if *folD* is required for cell-to-cell spread, we infected monolayers of rat fibroblasts with WT, Δ*folD* or Δ*folD*::*folD* Lm for 60 hours and measured the size of plaques formed by each strain. The area of plaques formed by Δ*folD* was nearly 70% smaller than that of WT, whereas the plaque size of Δ*folD*::*folD* bacteria was comparable with WT (Figure 3D). We saw similar but less drastic results when we measured plaque size in immortalized murine hepatocytes (Figure 3E). In contrast, Δ*folD* replicated to equal levels as WT in both naïve and IFN-γ-stimulated BMDMs (Figure S2C and S2D). In all, these data suggest that the requirement for folates produced by FolD, is cell-type specific and contributes to growth in the murine liver.

### Dysregulation of the *L. monocytogenes* D-allose utilization operon leads to decreased virulence in mice

D-allose is a C3 epimer of glucose which is found in low quantities in nature, primarily in plants (32). D-allose and its derivative D-allulose are gaining favor as safe, low-caloric sweeteners (32, 33). In addition, D-allose is used in a novel enrichment broth to preferentially culture pathogenic *Listeria* (*L. monocytogenes* and *L. ivanovii*) from environmental samples (34). A six gene operon *lmo0734-lmo0739* confers the ability for Lm to grow on D-allose as a sole carbon source (Figure 4A) (35). Thus, we were intrigued to find that the transcriptional regulator of this D-<u>al</u>lose <u>o</u>peron (ALO), *alsR* (*lmo0734*), was depleted over 70-fold in the liver and 45-fold in the spleen (Figures 2A and 2B and Tables S1 and S2). To validate the role of *alsR* in Lm *in vivo* pathogenesis, we used allelic exchange to make an in-frame deletion of *alsR,* infected WT C57BL/6 mice via IV, and enumerated CFU from livers and spleens 72 hpi. We observed a significant reduction in Δ*alsR* CFU in both livers (4.5 log_10_) and spleens (2.5 log_10_) compared to WT (Figures 4B and S3A). We complemented *alsR* back to a separate chromosomal locus with its native promotor (Δ*alsR*::*alsR*) and observed that these bacteria grew to the same levels as WT in both the liver and spleen (Figures 4B and S3A). To determine if Δ*alsR* bacteria were impaired prior to organ colonization, we repeated the IV infection but collected bacteria from livers and spleens at 4 and 25 hpi. At 4 hpi, we counted similar numbers of WT and Δ*alsR* CFU isolated from the livers of mice (Figure 4C) but observed a small decrease in Δ*alsR* bacteria isolated from spleens compared to WT (Figure S3B). In contrast, there were significantly fewer Δ*alsR* CFU in both organs at 25 hpi (Figures 4C and S3B). When observed over time, the growth of Δ*alsR* slowed after 4 hpi compared to WT (Figure S3C) and suggests that the Δ*alsR* growth defect occurs after organ colonization. Interestingly, although Δ*alsR* survival was impaired significantly *in vivo* as early as 25 hpi, we observed no difference in intracellular replication in BMDM between WT and Δ*alsR* (Figure S3D) and only a modest decrease in plaque size (Figure S3E).

**Figure 4.**
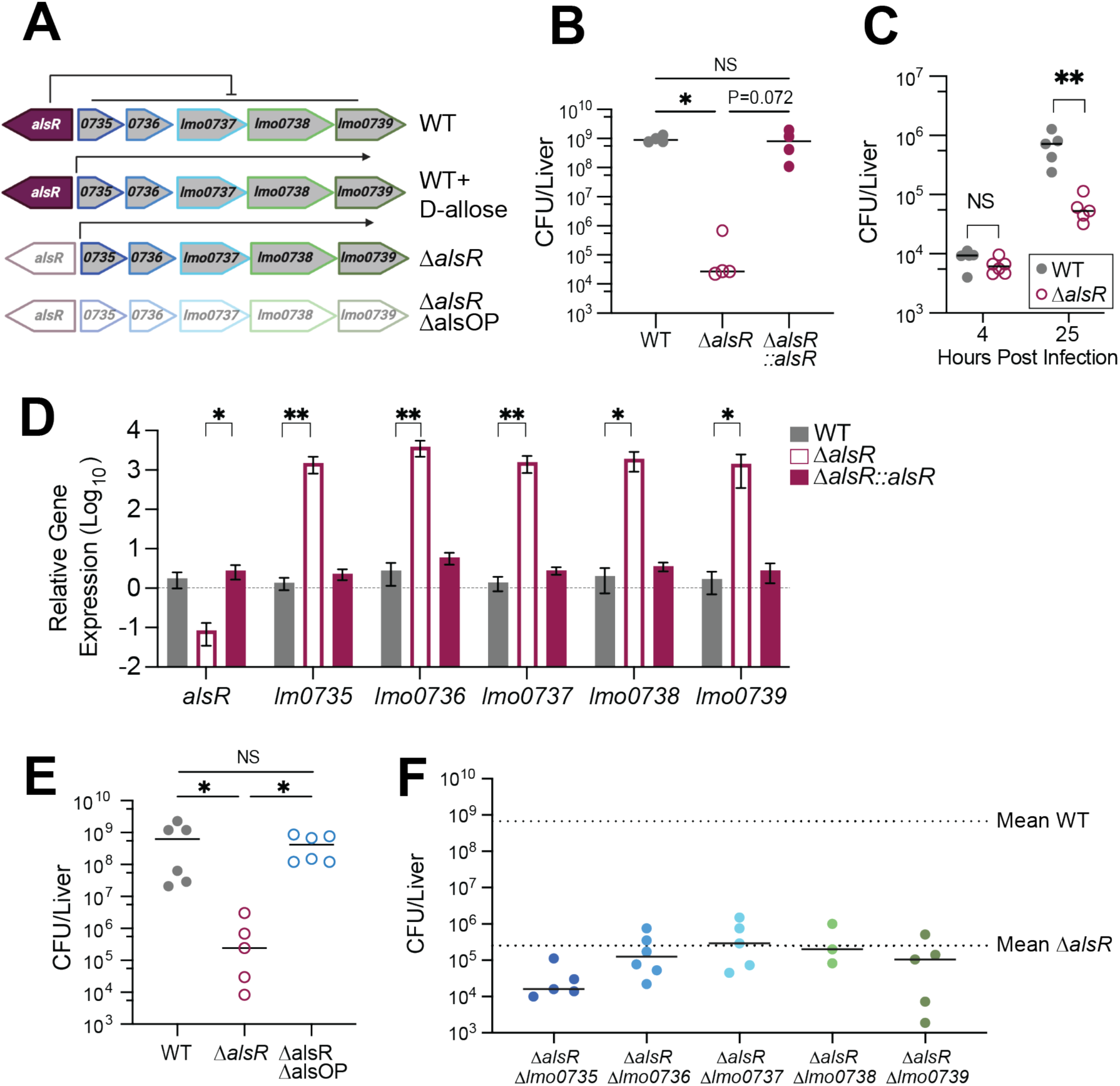
Dysregulation of the Lm D-allose operon leads to decreased virulence in mice. (**A**) Schematic of the ALO locus in Lm and expected expression under indicated conditions. (**B**) CFU harvested from livers of WT mice infected with WT, Δ*alsR* or Δ*alsR::alsR* Lm for 72 hours. (**C**) CFU harvested from livers of WT mice infected with WT or Δ*alsR* Lm for 4 or 25 hours. ** p<0.01 by Mann-Whitney analysis. (**D**) Relative gene expression of the ALO in WT, Δ*alsR* and Δ*alsR*::*alsR* Lm grown in BHI until mid-log phase. Data are plotted as fold change over WT for each gene. * P<0.05, ** P<0.01 by one way ANOVA of ΔΔC_t_ values. (**E**) WT mice were infected with WT, Δ*alsR* or Δ*alsR*ΔalsOP Lm and CFU were harvested from livers at 72 hpi. (**F**) WT mice were infected with the indicated strains and CFU were enumerated from livers at 72 hpi. Mean WT and Δ*alsR* CFU were calculated from at least four independent experiments. CFU for individual gene mutants in Δ*alsR* background were collected on independent days. Experiments in **C**, and **E-F** were performed once. CFU in **B** and **E** were analyzed by Kruskal-Wallis test with * p<0.05.

Because AlsR is a predicted repressor, we measured ALO expression in the mutant by qRT-PCR. We confirmed that in the absence of *alsR*, all genes in the operon were upregulated in BHI without the presence of D-allose (Figure 4D). Importantly, this overexpression did not result in *in vitro* growth defects in BHI compared to WT Lm (Figure S3F). Additionally, since phosphoenolpyruvate (PEP)-dependent phosphotransferase system (PTS) sugars suppress PrfA-dependent transcription (36), we explored whether aberrant expression of the ALO could influence virulence gene expression. To test this, we generated a strain of Δ*alsR* that contains a constitutively active mutant of PrfA (Δ*alsR* PrfA* (37)) and infected mice via IV. PrfA* did not rescue the Δ*alsR* growth defect in either livers or spleens of mice at 72 hpi (Figures S3G and S3H), suggesting Δ*alsR* does not affect virulence gene expression. Thus, because AlsR is not predicted to regulate other genes (RegPrecise) (38), we postulated that dysregulation of the ALO is the direct cause for the Δ*alsR in vivo* growth defect. To test this, we deleted the entire operon including *alsR* from the chromosome (Δ*alsR*ΔalsOP) and infected mice via IV. In contrast to Δ*alsR*, Δ*alsR*ΔalsOP bacteria replicated to WT levels in the liver and spleen at 72 hpi (Figures 4E and S3I), suggesting that dysregulation of the ALO is responsible for the impaired growth of Δ*alsR*. To determine if overexpression of a single gene in the ALO was sufficient to produce the Δ*alsR in vivo* growth defect, we deleted each gene individually in the Δ*alsR* background, performed IV infections of mice and enumerated CFU at 72 hpi. Fewer CFU were recovered from livers of mice infected with Δ*alsR*Δ*lmo0735* compared to Δ*alsR* alone (Figure 4F and Figure S3J). In contrast, Δ*alsR*Δ*lmo0736*, Δ*alsR*Δ*lmo0737*, Δ*alsR*Δ*lmo0738* and Δ*alsR*Δ*lmo0739* CFU were similar to Δ*alsR* CFU (Figure 4F and Figures S3K-N). In all, no single ALO gene deletion rescued the growth of Δ*alsR* to WT or Δ*alsR*ΔalsOP levels.

## Discussion

A genome-wide understanding of the Lm fitness determinants in the host has yet to be explored. Here, we generated a murine model, RECON^-/-^, that sustained high-dose, acute bacterial infection. We used this host model to perform an *in vivo* Tn-seq of the human pathogen Lm and successfully identified both previously known and novel virulence determinants.

Mice lacking RECON restricted Lm systemic infection and consequently were protected from Lm pathogenesis. We previously observed that loss of RECON augmented immune signaling via NFκB (22). This suggests that RECON^-/-^ mice are poised to respond quicker to bacterial infection which contributes to the bacterial restriction we observed in the context of Lm infection (Figures 1C-D and S1C). As c-di-AMP binding to RECON inhibits enzyme activity (22), the immune environment of the RECON^-/-^ mouse represents a c-di-AMP bound state and suggests that build-up of a RECON substrate influences cross-talk with the immune system and contributes to bacterial restriction. Prostaglandins and the α,β-unsaturated aldehydes produced from lipid peroxidation are intriguing substrate possibilities for RECON (39, 40). Indeed, 4-hydroxynonenal influences NF-κB-dependent transcription by covalently modifying an upstream ubiquitin-conjugating enzyme Ube2V1 (41). Work is ongoing to define the interaction of RECON (and its substrates or products) on the NF-κB signaling axis in RECON^-/-^ mice. However, we exploited the increased LD50 of the RECON^-/-^ mouse to perform *in vivo* Tn-seq of Lm. A third of the genes we identified were previously implicated in Lm virulence in the host (Tables S1 and S2), which confirms the usefulness of RECON^-/-^ as a screening model. However, we cannot rule out that some of the novel hits from our screen are specific to the predicted heightened immune response in our model.

We have completed a genome-wide screen for Lm *in vivo* survival. Excitingly, two thirds of the genes required for Lm growth in either livers or spleens of mice have not been implicated in virulence prior to our screen. This provides a wealth of novel genetic targets to explore for disease prevention and treatment. However, the Tn library we used was not saturated and short genes were likely underrepresented. In fact, the first three genes of the ALO (*lmo0735-lmo0737*) were not present in our input library despite the expectation that they would not be essential in rich broth. This suggests that there may be more genes important for *in vivo* infection still to be discovered. In addition, the analysis we performed only mapped open reading frames (ORF) to the reference genome and did not take into account placement of small, non-coding RNAs (ncRNA) which are known to influence Lm virulence (42). For example, the multicopy ncRNA *lhrC5* is positioned downstream of *lmo0947*, which was depleted in our screen (43). Deletion of all five *lhrC* ncRNAs decreases Lm survival in macrophages (44). It is possible that Tn disruption of *lmo0974*, or similarly juxtaposed ORF-ncRNA pairs, decreased Lm survival due to altered ncRNA expression and subsequent target regulation. Finally, 9% of the genes required for Lm *in vivo* growth are unannotated. Further characterization of this unannotated subset is needed to understand the function of these genes in Lm physiology and their possible role as host virulence determinants.

Recently a Tn-seq was performed for Lm in J774 macrophages (10). Our screen identified nearly half (20/42) of the genes required for growth in macrophages which shows our methods were complementary. The lack of a total overlap between the two datasets is largely a result of our library density, as previously mentioned. For the remaining 22 genes required for macrophage growth, six were not represented in our input library and ten were also depleted in our screen but did not reach statistical significance. However, our screening *in vivo* led to identification of a larger subset of genes (139 compared to 42). One explanation for this is that many single gene deletions do not have a macrophage growth defect. Thus, screening *in vivo* allowed us to identify factors such as the known virulence genes *actA* (11) *gshF* (45) and *iap* (p60) (46), the lysosome resistance ABC transporter *eslAB* (47), the redox-responsive regulator *rex* (48) and alsR, which are all absent from the J774 dataset. In addition, although both screening conditions highlighted the requirement for lipoic acid uptake, and *de novo* purine and menaquinone synthesis, other nutrient restrictions only became apparent through screening *in vivo*. For example, the uptake genes for thiamine (*lmo1429*) and biotin (*lmo0598*) were only required in our *in vivo* screen despite being required for host infection (49, 50). Thus, by screening *in vivo* we identified genes involved in pathways that cell culture cannot recapitulate.

We separately characterized genes required for Lm survival in murine spleens and livers and identified 90 genes uniquely required for growth in the liver. It is likely that this number is an overrepresentation considering over 30 of the genes were also depleted in spleens at least two-fold but did not reach statistical significance. For some, such as the flavin metabolism gene *ribF*, and a menaquinone synthesis gene *menA*, the p-value was just below our cutoff of p<0.05, emphasizing the requirement of these essential cofactors for *in vivo* growth. Interestingly, while glycerol utilization genes (ex. *pgm*, and *pfkA*) were required in both organs, there seems to be a liver specific energy requirement not found in the spleen. The first indication for this is that nine membrane transport mechanisms were required in the liver only, including several genes, *pstB* and its regulator *phoU*, *lmo1849,* and *lmo0366* for ion transport. In addition, we also identified several genes for ethanolamine usage (*eutC*, *eutE* and *lmo1161*) which can contribute to acetyl-coA production (51) or act as a nitrogen source (52) from host phosphatidylethanolamine. Finally, we characterized the liver-specific requirement for the folate enzyme FolD which was recently described as an N-formylmethionine-dependent phenotype by Feng and colleagues (53). We observed a larger growth defect for Δ*folD* in livers, likely because we used a different mouse strain and harvested CFU at a later time point. That we independently identified *folD* and observed similar phenotypes again emphasizes the universal benefit of the RECON^-/-^ model for bacterial fitness evaluation.

The D-allose utilization repressor gene *alsR* was required for Lm survival in both livers and spleens of mice but not in *in vitro* cell culture models. Overexpression of the ALO in the Δ*alsR* background was sufficient to cause the *in vivo* growth defect, which reinforces the importance of tight metabolic regulation in the host (54, 55). Indeed, the loss of two other regulators, *lmo0020* and *lmo1253* that control PTS^Man^-1 and trehalose metabolism, respectively, also impaired Lm survival in the liver (Table S1). However, expression of the ALO in the environment likely benefits Lm to outcompete other microbes in the community and this growth advantage is exploited to enrich Lm during sampling studies (34). Interestingly, the ALO is only encoded by Lineage II Lm and is used to distinguish Lineage II from Lineage I and III strains in PCR-based serotyping methods (56). Since we were able to delete the ALO and observed WT growth in the host, it is not surprising that Lineage I strains, which are frequently associated with human Listeriosis outbreaks (4), have lost the operon. In contrast, some Lineage I serotypes have acquired the ALO (57, 58) which not only calls into question the accuracy of using the ALO as a serotype marker (59) but also suggests that acquisition of the operon is beneficial to Lm prior to human colonization, such as during contamination of food processing environments. It is intriguing to speculate how the Δ*alsR* phenotype could be utilized to prevent disease. Since AlsR is predicted to relieve repression upon binding D-allose, identification of a non-convertible, synthetic activator would lead to expression of the ALO without the growth advantage provided by the endogenous substrate, D-allose. Thus, conceivably, addition of the synthetic activator to ready to eat foods could act as a Listeriosis prophylactic.

## Methods

### Bacterial Growth Conditions

All *L. monocytogenes* strains used in this study were on the background of the WT strain 10403S. Lm was grown in BHI broth or on BHI+1.5% agar (RPI # B11000-5000.0).

Overnight cultures of Lm were obtained by growth at 30°C statically for 18 h. When needed, cultures were supplemented with the following antibiotics: chloramphenicol (5-10 µg/mL), erythromycin (2 µg/mL), streptomycin (200 µg/mL), or tetracyline (2 µg/mL). Unless otherwise specified, catalog numbers refer to Thermo Scientific products. Donor *E. coli* SM10 were grown in LB broth (Fisher Scientific BP97235) or LB+1.5% agar supplemented when needed with carbenicillin (100 µg/mL) or chloramphenicol (34 µg/mL).

### Plasmid and Strain Construction

The strains used in this study are listed in Table S3 and primers in Table S4. Chromosomal deletions were made using allelic exchange from the pLIM backbone followed by counterselection on 18 mM DL-4-chlorohenylalanine (#157280250) (60). Complementation strains were made by integrating pPL2 (61) to the chromosome under pHyper (62) (*folD*::*folD*) or the native promoter (*alsR*::*alsR*) with selection on tetracyline. All plasmids were introduced to Lm via conjugation from *E. coli* SM10. A Lm transposon library was constructed as previously described (63) and stored at -80°C in 1 mL aliquots in BHI+ 40% glycerol.

### Mice

Animal experiments strictly complied with the recommendations outlined in the Guide for the Care and Use of Laboratory Animals by the National Research Council. Experiments were performed under protocols reviewed and approved by the University of Washington Institutional Animal Care and Use Committee. Animals were housed at the University of Washington Department of Comparative Medicine vivarium under specific pathogen free conditions. The heterozygous RECON^+/-^ founder as well as age-matched wildtype controls were bred with mice purchased from Jackson Labs (#000664).

### Generation of RECON-deficient mice

CRISPR/Cas9-engineered mice were generated as previously described (64, 65) with the University of Washington Transgenic Resources Program. Guide RNAs targeting exon 6 of *Akr1c13* were cloned into pX330-U6-Chimeric_BB-CBh-hSpCas9 (Addgene #42230). A T7-sgRNA PCR product was amplified and *in vitro* transcribed as previously described (65) using the MEGAshortscript T7 kit (#AM1354) and purified with the MEGAclear kit (#AM1908). *Cas9* mRNA used for injections was purchased from Sigma (#CAS9MRNA). Primer sequences are provided in the Table S4.

### RGEN-RFLP assay

Genotyping of founder mice using CRISPR/Cas-derived RNA-guided engineered nucleases (RGEN) restriction fragment length polymorphism (RFLP) analysis was done as previously described (64, 66). Guide RNAs were cloned and purified as detailed above. A 1.2kb amplicon spanning the *Akr1c13* exon 6 genomic region was used as substrate DNA. Primer sequences are provided in Table S4. The RGEN-RFLP assay with Cas9 nuclease from *S. pyrogenes* was carried out according to manufacturer’s instructions (NEB #M0386) with 30 nM sgRNA and 3 nM substrate DNA final concentrations.

### Primary Macrophages

Primary BMDM from C57BL/6 and RECON^-/-^ mice were isolated (67) and differentiated (68) as previously described. BMDM were grown at 37°C in 5% CO_2_ in Dulbecco’s Modified Eagle Medium (DMEM) GlutaMAX (Gibco #10569-010) supplemented with 1 mM sodium pyruvate, 55 µM 2-mercaptoethanol, 10% heat-inactivated FBS and 10% L929 conditioned medium.

### RNA isolation and qRT-PCR analysis

For *Akr1c13* expression in tissues, organs were harvested, placed in 1mL LBP from the NucleoSpin RNA Plus kit (Clontech #740984.250) with silica disruption beads and snap frozen in liquid nitrogen. Samples were thawed on ice, homogenized with a multivortexer, and RNA was isolated with NucleoSpin RNA Plus kit per manufacturer’s instructions. cDNA was synthesized using Maxima H Minus First Strand cDNA synthesis kit (#EP0752). TaqMan Gene Expression Assay probes were used for quantification of gene expression (#Mm00657347_m1) with *Hprt* (#Mm03024075_m1) as an endogenous control. For ALO gene expression, overnight Lm were diluted to OD_600_=0.05 in BHI and grown at 37°C with shaking until OD_600_=0.4-0.6 and RNA was extracted as previously described (69). Briefly, equal volumes of culture and ice cold 100% methanol were mixed on ice and bacteria were pelleted, flash frozen and stored at -80°C. RNA was extracted using acidified phenol:chloroform:isoamyl alcohol with vortex agitation. RNA was precipitated, DNAse treated (#AM1907) and reverse transcribed using iScript cDNA synthesis (Bio-Rad #1708891). qPCR was performed using SYBR Green Master Mix (#A25742). Primers for qPCR are listed in Table S4.

### Mouse infections

All infections were carried out in 7-12 week old RECON^-/-^ or C57BL/6 mice with equal sex distribution. For Tn-seq, a single vial of a Lm transposon library of approximately 17,000 mutants was thawed, diluted in 3mL BHI, grown three hours at 37°C, 240 rpm. The culture was then diluted in sterile phosphate buffered saline (PBS) to 1.2 x 10^7^ CFU/mL and 200 µL of this suspension was administered IV through the retroorbital injection to RECON^-/-^ mice to give 2.4 x 10^6^ CFU/mouse. Mice were sacrificed at 34 hpi and total liver and spleen homogenates were plated on BHI agar. Bacterial biomass was scraped and stored at -80°C. For all other infections, Lm were prepared as follows: overnight cultures of Lm were back-diluted in BHI (1.2 mL into 4.8 mL) and grown for 1 h at 37°C shaking then diluted in sterile PBS accordingly. For mouse survival studies, mice were IP injected with 1×10^6^ CFU/mouse in 200 µL. For experiments enumerating tissue CFU from RECON^-/-^, 1×10^4^ CFU/mouse was injected IP in 200 µL and 72 hpi the livers and spleen were collected, homogenized in 0.1% IGEPAL, and plated for CFUs. All remaining infections were carried out in C56BL/6 mice with 1×10^5^ Lm CFU delivered IV in 200 µL. At indicated time points, livers and spleens were collected and homogenized in 10 mL or 5 mL (respectively) 0.1% IGEPAL and plated on BHI agar with streptomycin.

### Plaque Assay

Immortalized L2 rat fibroblasts and TIB73 mouse hepatocytes were cultures in DMEM GlutaMAX with 1mM sodium pyruvate and 10% FBS. Plaque assays were performed on monolayers of L2 or TIB73 cells as previously described (70) with the following modifications. Cells were seeded in 3 mL media/well. Overnight Lm cultures were pelleted and resuspended in 1X PBS to OD_600_=1.0, then diluted 1:30 in PBS. To infect, 5 µL of the diluted bacteria were added directly to each well and swirled to mix (MOI 0.2). After one hour, cells were washed twice in 1X PBS prior to the addition of an agar plug (DMEM+10% FBS, 1 mM sodium pyruvate, 2 mM glutamine, 0.7% agarose, 10 µg/mL gentamicin). After 2 days, the staining mix (DMEM+10% FBS, 1 mM sodium pyruvate, 2 mM glutamine, 0.7% agarose, 0.25% Neutral Red) was added for 18 h before plates were scanned and plaques analyzed in ImageJ. Superpure agarose was purchased from Biotech Sources (#G02PD-125) and Neutral Red from Sigma (#N6264).

### Tn-seq Library Construction and Sequencing

Genomic DNA was extracted using MasterPure Gram-Positive DNA Purification Kit (Fisher Scientific #NC9197506) with 300 U/mL mutanolysin (Sigma #M9901-10KU) in place of lysosome. DNA was diluted to 3 µg/130 µL in microTUBEs (Covaris #520045) and sheared in triplicate to 300 bp fragments using the following settings on a Covaris LE220 Focused-Ultrasonicator: duty cycle 10%, peak intensity 450, cycles per burst 100, duration 200 s/column. Fragmented DNA was end-repaired (NEB #E6050) and purified on Ampure SPRIselect beads (Beckman-Coulter #B23317). Poly-C-tails were added to 1 µg of each end-repaired sample in duplicate using Terminal Transferase (Promega) with a ratio of 9.5 mM dCTP to 0.5 mM ddCTP to limit chain length and DNA Duplicate reactions were combined and purified with SPRIselect beads. Transposon junctions were amplified with oligos pJZ_Fwd_RND1 and olj376 using 500 ng DNA and KAPA HiFi Hotstart Mix (Kapa Biosystems, #KK2602). Reactions were stopped at the inflection point of amplification (6-14 cycles). Transposon junction amplicons were purified using SPRIselect beads. Finally, barcoded adaptors were added to using KAPA HiFi Hotstart Mix, pJZ_Fwd_RND2 and adaptors for pooled sequencing (Table S4). The amount of each sample to add was empirically determined such that each sample reached inflection point after 17 rounds. DNA was purified and size selected on SPRIselect beads for fragments 250-450 bp in size. Samples were pooled to 9 nM and sequenced as single end 50 bp reads on NextSeq HO with a 10% PhiX spike in.

### Conditional Essentiality and Pathway Analysis

The *L. monocytogenes* 10403S NC_017544 genome was uploaded to PATRIC (24) as the reference genome. Trimmed reads were mapped and assessed for essentiality using TRANSIT with resampling (71). All genes that reached a cut off of p<0.05 with at least a two-fold depletion were considered essential *in vivo*. These genes were further analyzed for KEGG Pathway enrichment using DAVID (72).

## Data Availability

Tn-seq data generated in this study have been deposited to the Sequence Reads Archive under BioProject PRJNA1062841 (http://www.ncbi.nlm.nih.gov/bioproject/1062841) and will be publicly available at the date of publication.

## Stats and Quantification

All numerical data were analyzed and visualized in GraphPad Prism 9.0 software. For plaque assay and murine infection data, all data points are plotted with the bar at the median. The ALO qRT-PCR data are represented as mean with standard deviation. The number of repeats and statistical test used for each experiment is detailed in the figure legends.

## Acknowledgements

We would like to thank Larry Gallagher and Hannah Ledvina for help with library preparation design, and Susan Brewer, Ena Chen, and Lieselotte Kreuk for experimental support. Additionally, we thank members of the Reniere lab at the University of Washington (UW) for valuable discussion and feedback. We thank the Mougous, Reniere and Smith labs at UW for reagents and equipment sharing. Schematics were created with Biorender.com. A.P.M was supported by the National Science Foundation Graduate Research Fellowship Program (#DGE-1256082), M.L.N was a Cancer Research Institute Irvington Fellow supported by the Cancer Research Institute (CRI Award #CRI3450) and H.T was supported by a National Institute of General Medical Sciences Public Health Service National Research Service Award (#T32GM007270). This work was also supported by National Institute of Allergy and Infectious Diseases R01AI116669 (J.J.W).

---

**Supplemental Figure 1.**
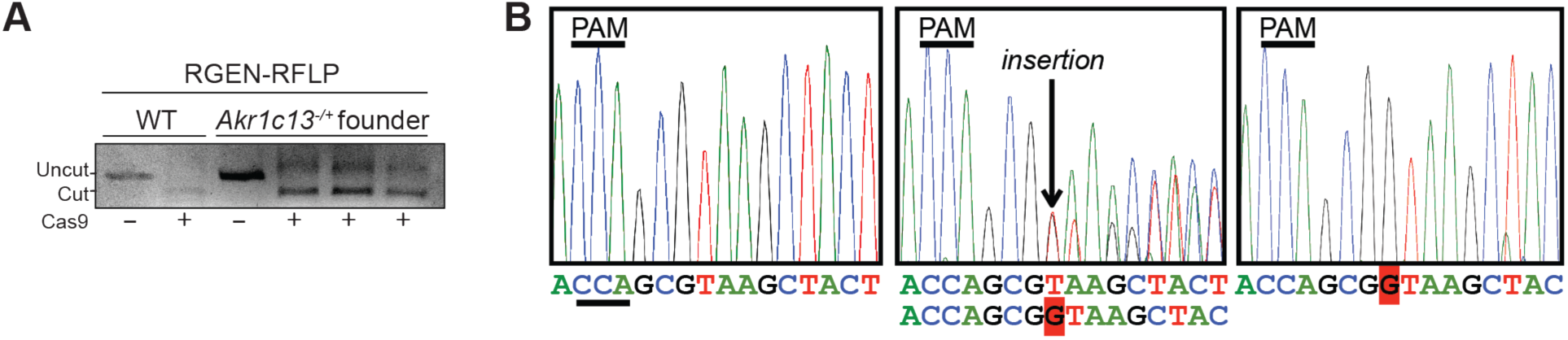
Generation of RECON-deficient mice using CRISPR/Cas9-mediated mutagenesis. (**A**) RGEN-RFLP analysis of CRISPR-targeted region (exon 6) in the *Akr1c13* gene in the heterozygous founder. (**B**) Sequence of mutated allele with a single G insertion leading to a frameshift mutation.

**Supplemental Figure 2.**
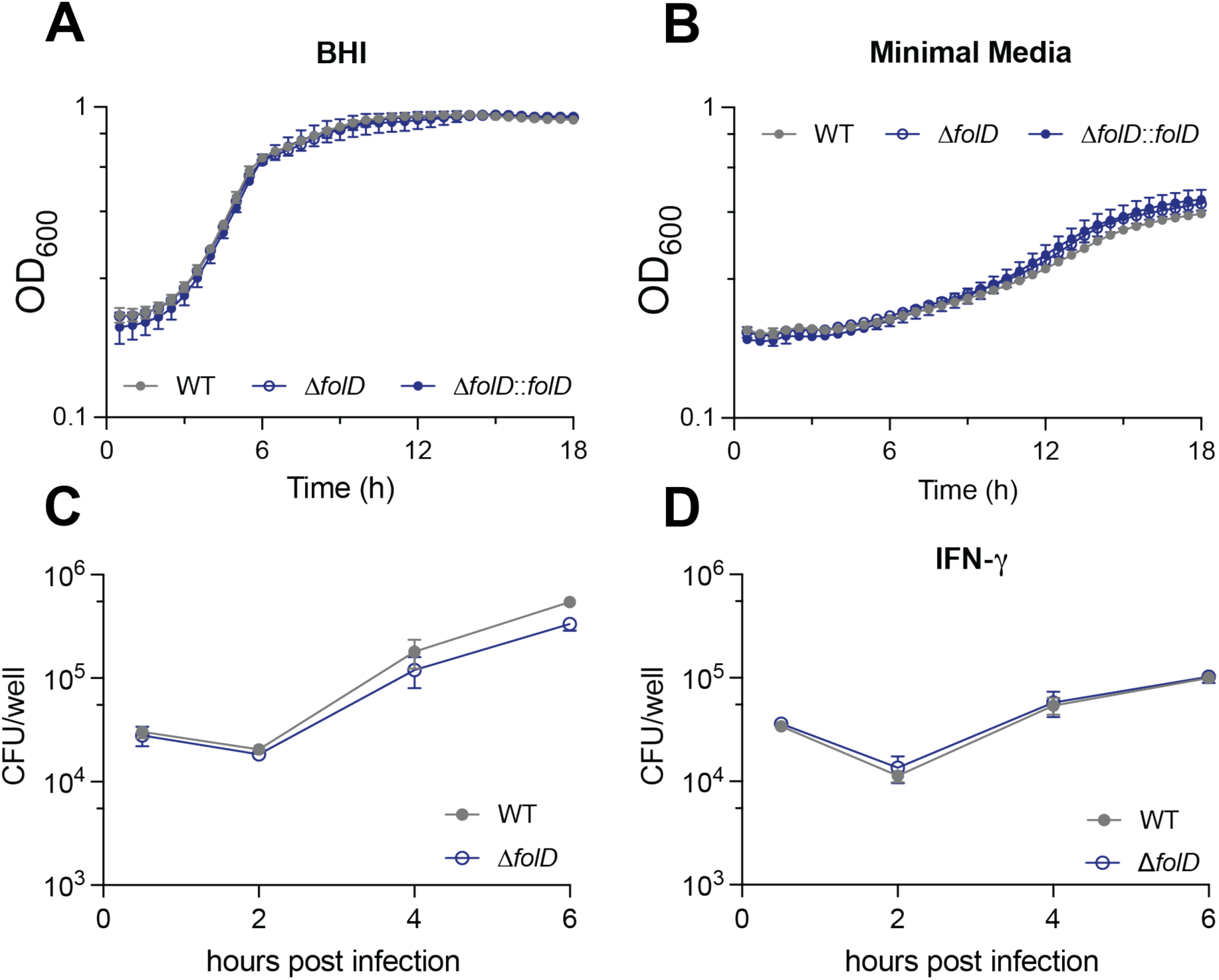
The folate cycle gene *folD* is not required for Lm *in vitro* growth or growth in macrophages. (**A-B**) Bacterial growth as measured by OD600 over time in BHI (**A**) or minimal media (**B**). (**C-D**) BMDMs were left naïve (**C**) or stimulated with IFN-γ (**D**) for 18 hours prior to infection with Lm. Intracellular CFU were collected at the indicated time points.

**Supplemental Figure 3.**
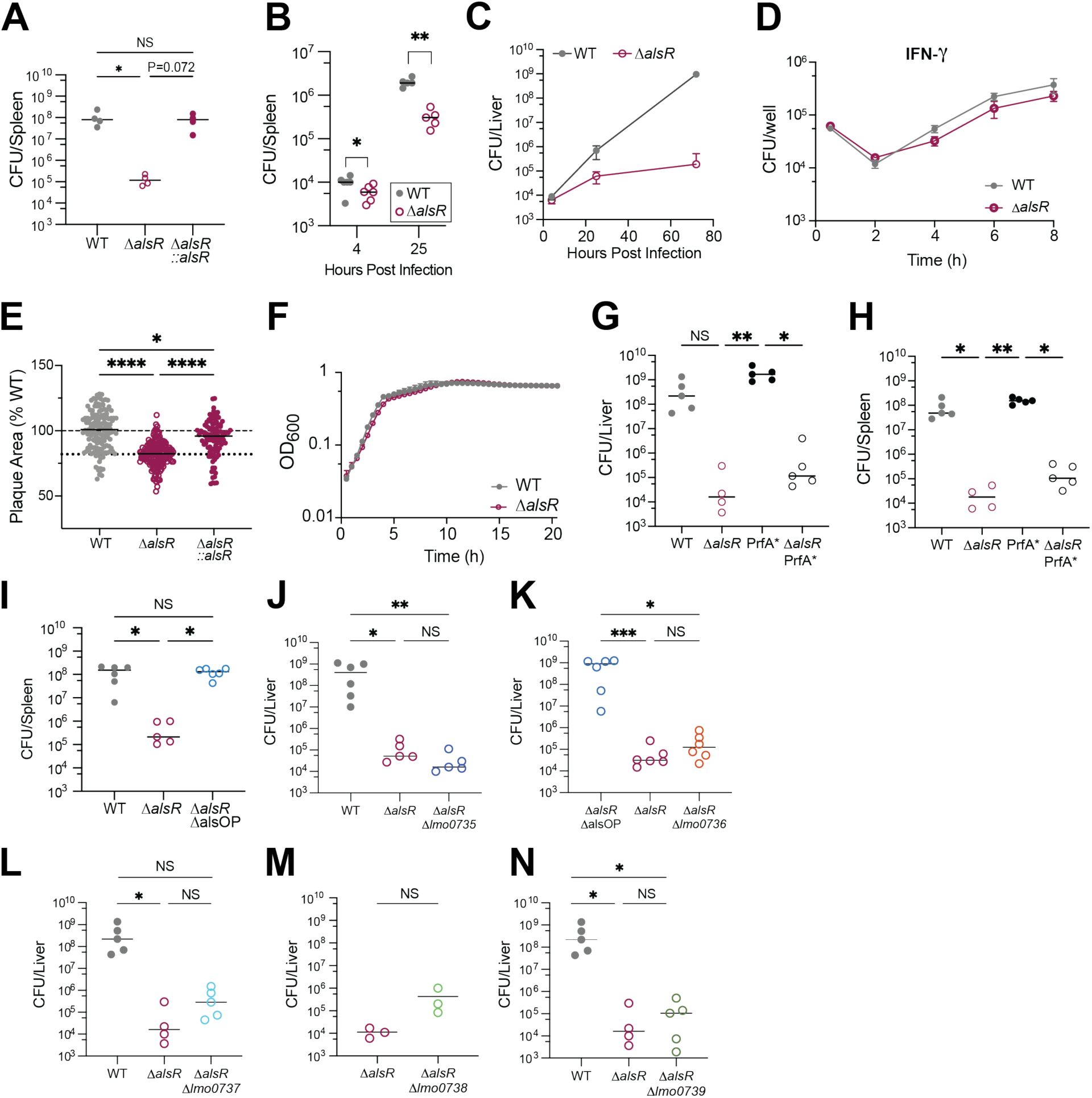
Dysregulation of the D-allose utilization operon impairs Lm growth *in vivo* but not *in vitro*. (**A**) CFU harvested from spleens of WT mice infected with WT, Δ*alsR* or Δ*alsR::alsR* Lm for 72 hours. (**B**) CFU harvested from spleens of WT mice infected with WT or Δ*alsR* for 4 or 25 hours. (**C**) Mean CFU of WT or Δ*alsR* from panels Figures 4B and 4C plotted over time. (**D**) IFN-γ-stimulated BMDMs were infected with Lm for the indicated time points and intracellular CFU were enumerated. (**E**) Plaque area measured in fibroblasts infected with WT, Δ*alsR* or Δ*alsR*::*alsR* Lm for 48 hours and stained with neutral red. (**F**) Bacterial growth as measured by OD_600_ over time in BHI. (**G-H**) CFU harvested from livers (**G**) and spleens (**H**) of WT mice infected with WT, Δ*alsR,* PrfA* or Δ*alsR* PrfA* Lm for 72 hours. (**I**). CFU harvested from spleens of WT mice infected with WT, Δ*alsR* or Δ*alsR*ΔalsOP Lm for 72 hours. (**J-N**) CFU harvested from livers of WT mice infected with the indicated strains for 72 hours. Experiments in **B**, **G-H**, and **I-N** were performed once. Plaque area in **E** and CFU in **A, G-N** were analyzed by Kruskal-Wallis test with * p<0.05, ** p< 0.005 and **** p< 0.0001. CFU in **B** was analyzed by Mann-Whitney analysis with *p<0.05, **p<0.01.

**Supplemental Table 1.**
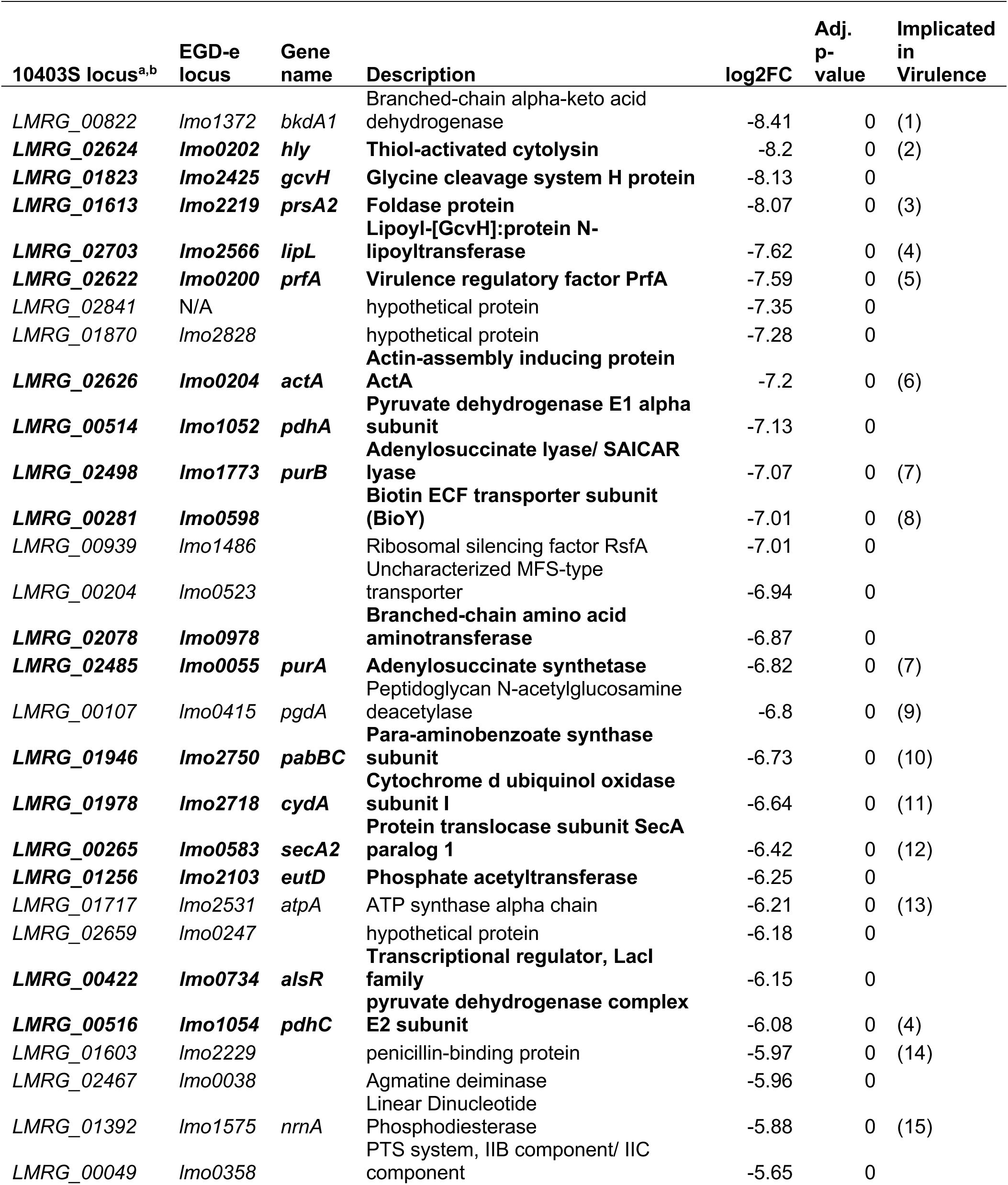
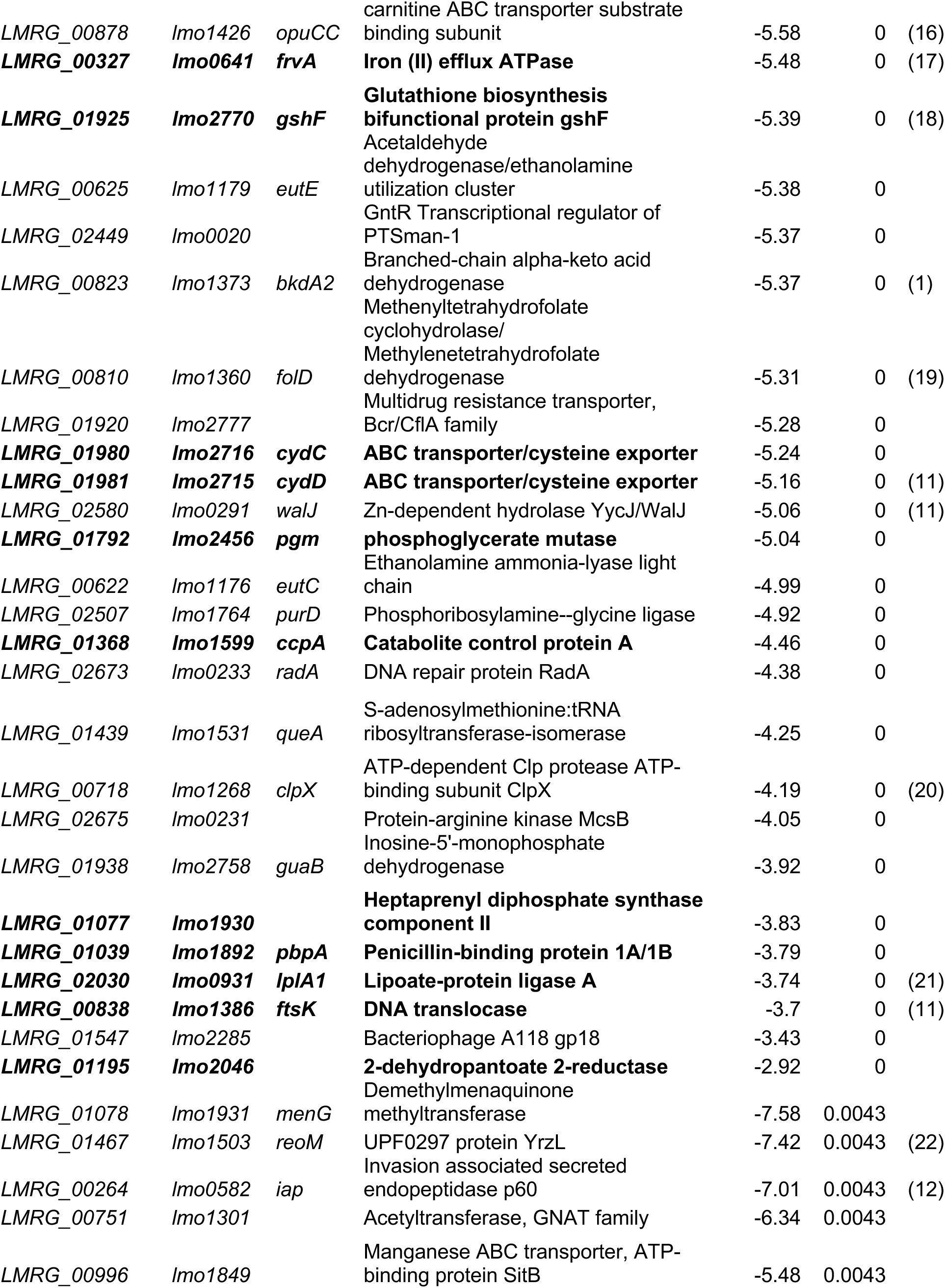
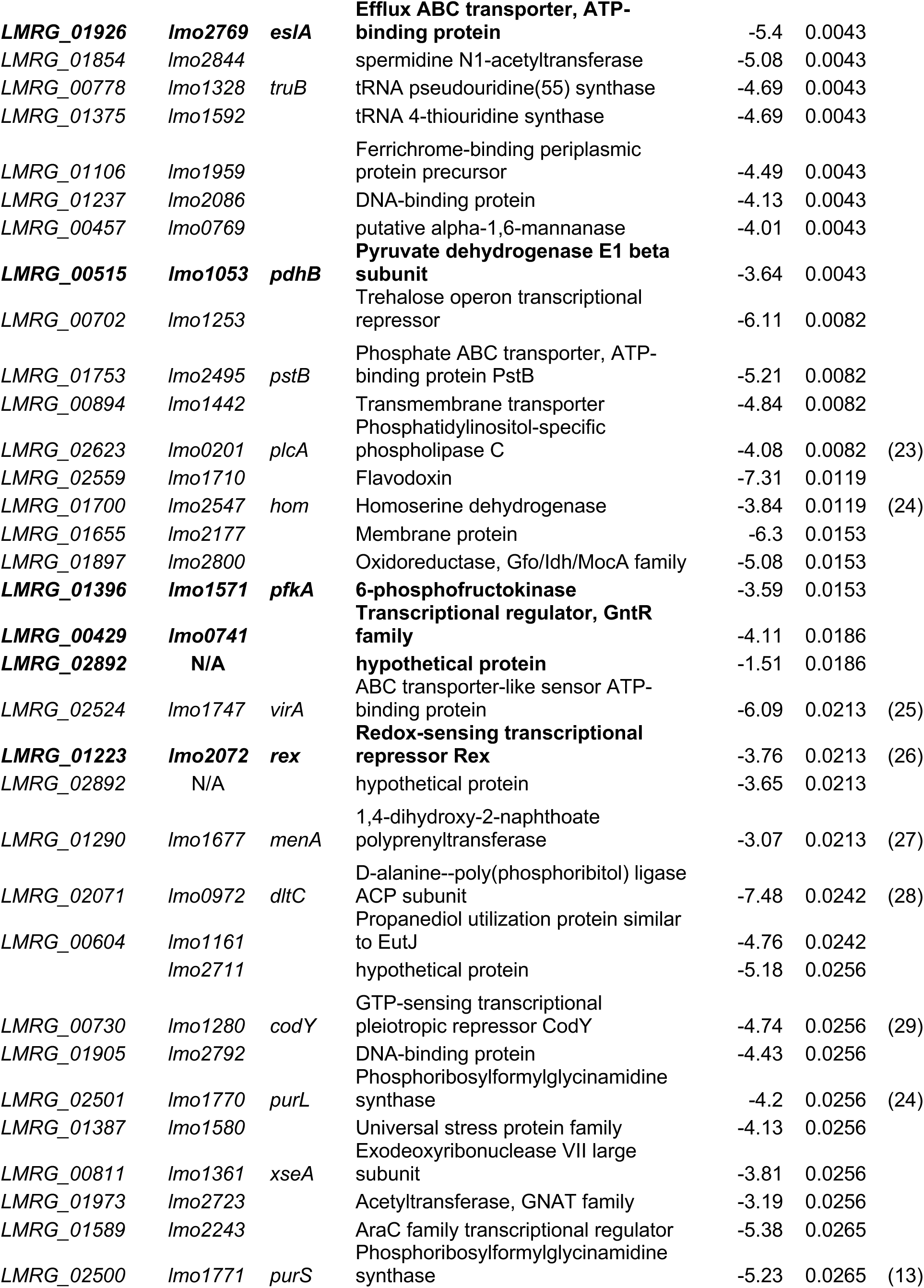
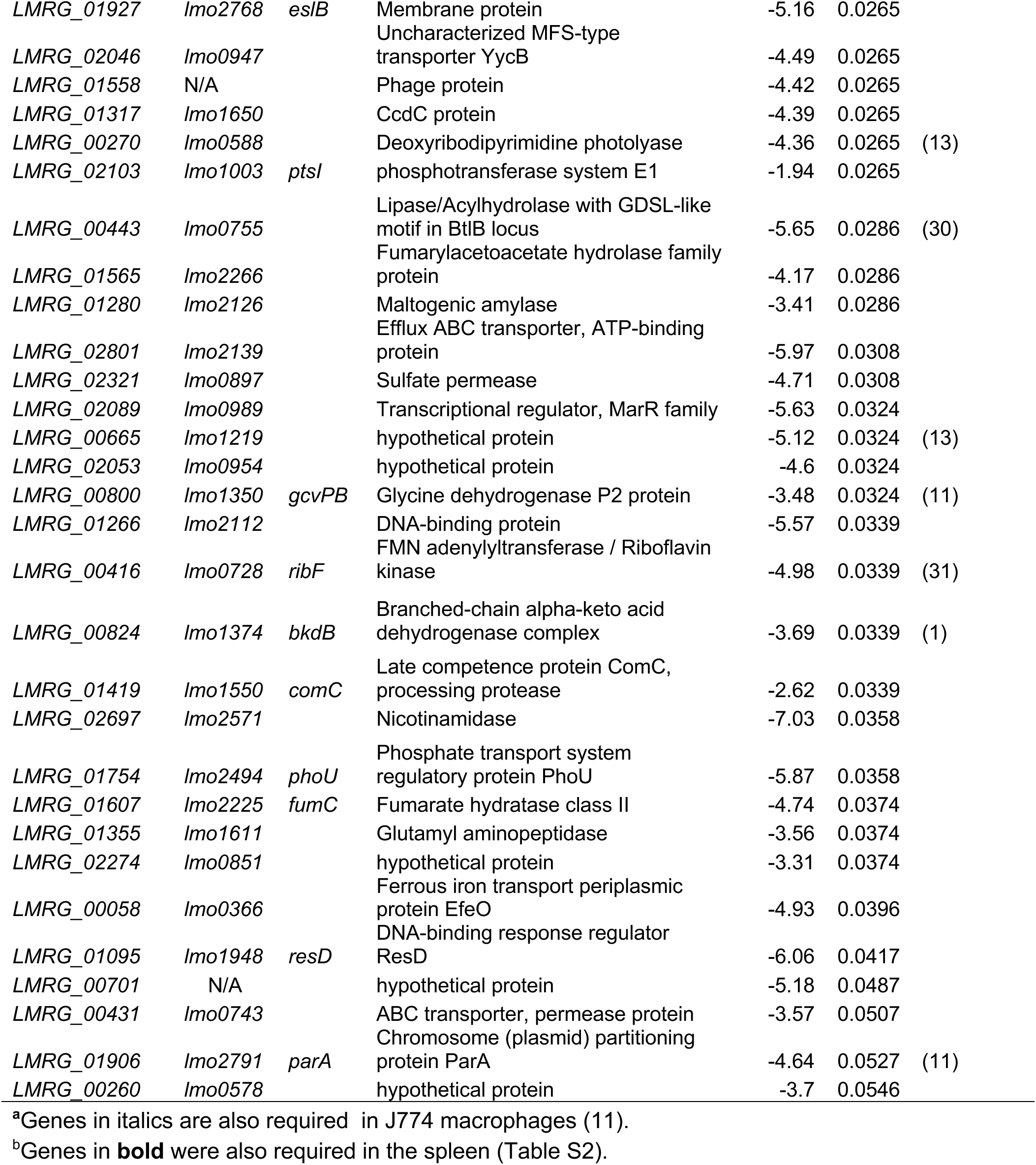
Lm genes required for survival in the murine liver.

**Supplemental Table 2.** Lm genes required for survival in the murine spleen.

| 10403S Locus<sup>a,b</sup> | EGD-e locus | Gene Name | Description | log2FC | Adj. p-value | Implicated in virulence |
| --- | --- | --- | --- | --- | --- | --- |
| *LMRG_01823* | *lmo2425* | *gcvH* | Glycine cleavage system H protein | -8.43 | 0 |  |
| *LMRG_01925* | *lmo2770* | *gshF* | Glutathione biosynthesis bifunctional protein gshF | -8.36 | 0 | (18) |
| *LMRG_02624* | *lmo0202* | *hly* | Thiol-activated cytolysin | -8.13 | 0 | (2) |
| *LMRG_01613* | *lmo2219* | *prsA2* | Foldase protein | -7.94 | 0 | (3) |
| *LMRG_02622* | *lmo0200* | *prfA* | Virulence regulatory factor PrfA | -7.9 | 0 | (5) |
| *LMRG_01774* | *lmo2474* |  | RNase adapter protein RapZ | -7.66 | 0 |  |
| *LMRG_02703* | *lmo2566* | *lipL* | Lipoyl-[GcvH]:protein N-lipoyltransferase | -7.62 | 0 | (4) |
| *LMRG_00514* | *lmo1052* | *pdhA* | Pyruvate dehydrogenase E1 component alpha subunit | -7.39 | 0 |  |
| *LMRG_02498* | *lmo1773* | *purB* | Adenylosuccinate lyase SAICAR lyase | -6.88 | 0 | (7) |
| *LMRG_00281* | *lmo0598* |  | Biotin ECF transporter subunit (BioY) | -6.88 | 0 | (8) |
| *LMRG_02718* | *lmo2718* | *cydA* | Cytochrome d ubiquinol oxidase subunit I | -6.65 | 0 | (11) |
| *LMRG_00516* | *lmo1054* | *pdhC* | pyruvate dehydrogenase complex E2 subunit | -6.41 | 0 | (4) |
| *LMRG_01926* | *lmo2769* | *eslA* | Efflux ABC transporter, ATP-binding protein | -6.34 | 0 |  |
| *LMRG_02485* | *lmo0055* | *purA* | Adenylosuccinate synthetase | -6.03 | 0 | (7) |
| *LMRG_00265* | *lmo0583* | *secA2* | Protein translocase subunit SecA paralog 1 | -6.02 | 0 | (12) |
| *LMRG_01256* | *lmo2103* | *eutD* | Phosphate acetyltransferase | -5.66 | 0 |  |
| *LMRG_00422* | *lmo0734* | *alsR* | Transcriptional regulator, LacI family | -5.51 | 0 |  |
| *LMRG_00327* | *lmo0641* | *frvA* | Iron (II) efflux ATPase | -5.46 | 0 |  |
| *LMRG_01980* | *lmo2716* | *cydC* | ABC transporter/cysteine exporter | -5.21 | 0 |  |
| *LMRG_01981* | *lmo2715* | *cydD* | ABC transporter/cysteine exporter | -5.08 | 0 | (11) |
| *LMRG_02030* | *lmo0931* | *lplA1* | Lipoate-protein ligase A | -4.94 | 0 | (21) |
| *LMRG_01792* | *lmo2456* | *pgm* | phosphoglycerate mutase | -4.72 | 0 |  |
| *LMRG_01946* | *lmo2750* | *pabBC* | Para-aminobenzoate synthase subunit I | -4.39 | 0 | (10) |
| *LMRG_02460* | *lmo0031* |  | Transcriptional regulator, LacI family | -4.21 | 0 |  |
| *LMRG_01223* | *lmo2072* | *rex* | Redox-sensing transcriptional repressor Rex | -4.17 | 0 | (26) |
| *LMRG_01077* | *lmo1930* |  | Heptaprenyl diphosphate synthase component II | -4 | 0 |  |
| *LMRG_00515* | *lmo1053* | *pdhB* | Pyruvate dehydrogenase E1 beta subunit | -3.71 | 0 |  |
| *LMRG_00838* | *lmo1386* | *ftsK* | DNA translocase FtsK | -3.7 | 0 | (11) |
| *LMRG_02627* | *lmo0205* | *plcB* | Broad-substrate range phospholipase C | -3.52 | 0 | (32) |
| ***LMRG_01195*** | ***lmo2046*** |  | **2-dehydropantoate 2-reductase** | -3.28 | 0 |  |
| ***LMRG_01368*** | ***lmo1599*** | ***ccpA*** | **Catabolite control protein A** | -3.22 | 0 |  |
| *LMRG_02636* | *lmo0214* | *mfd* | Transcription-repair coupling factor | -2.1 | 0 |  |
| ***LMRG_02626*** | ***lmo0204*** | ***actA*** | **Actin-assembly inducing protein ActA** | -6.95 | 0.0084 | (6) |
| *LMRG_01386* | *lmo1581* | *ackA* | Acetate kinase | -3.84 | 0.0084 |  |
| ***LMRG_01039*** | ***lmo1892*** | ***pbpA*** | **Penicillin-binding protein 1A/1B** | -3.03 | 0.0084 |  |
| *LMRG_02063* | *lmo0964* |  | Thioredoxin-fold protein | -5.11 | 0.0159 |  |
| *LMRG_01701* | *lmo2546* | *thrC* | Threonine synthase | -2.89 | 0.0159 |  |
| ***LMRG_02078*** | ***lmo0978*** |  | **Branched-chain amino acid aminotransferase** | -2.69 | 0.0232 |  |
| *LMRG_01923* | *lmo2772* |  | PTS system, beta-glucoside-specific | -1.91 | 0.0302 |  |
| *LMRG_00881* | *lmo1429* | *thiT* | Thiamine ECF transporter | -4.71 | 0.0359 | (33) |
| *LMRG_01204* | *lmo2054* |  | UPF0298 protein YlbG | -3.46 | 0.0359 |  |
| ***LMRG_00429*** | ***lmo0741*** |  | **Transcriptional regulator, GntR family** | -4.19 | 0.0468 |  |
| ***LMRG_01396*** | ***lmo1571*** | ***pfkA*** | **6-phosphofructokinase** | -3.57 | 0.0468 |  |
| *LMRG_02625* | *lmo0203* | *mpl* | Zinc metalloproteinase aureolysin | -2.97 | 0.0468 | (34) |
| ***LMRG_02892*** | **N/A** |  | **hypothetical protein** | -1.51 | 0.0523 |  |

<sup>a</sup>Genes in *italics* are also required in J774 macrophages (11).
<sup>b</sup>Genes in **bold** were also required in the liver (Table S1).

**Supplemental Table 3.** Strains used in this study.

| Strain Name | Background | Genotype | Plasmid | Reference |
| --- | --- | --- | --- | --- |
| JW655 | *E. coli* XL1Blue |  | pLIM | (35) |
| CS207 | *E. coli* SM10 | | pLIM-$\Delta folD$ | This study |
| CS208 | *E. coli* SM10 | | pLIM-$\Delta alsR$ | This study |
| CS210 | *E. coli* SM10 |  | pPL2x-pHyper-*folD* | This study |
| CS211 | *E. coli* SM10 |  | pPL2t-n200-*alsR* | This study |
| CS212 | *E. coli* SM10 | | pLIM-$\Delta alsR\Delta alsOP$ | This study |
| CS102 | *L. monocytogenes* 10403S |  |  | (36) |
| CS141 | *L. monocytogenes* 10403S | $\Delta folD$ | | This study |
| CS148 | *L. monocytogenes* 10403S | $\Delta folD::folD$ | | This study |
| CS147 | *L. monocytogenes* 10403S | $\Delta alsR$ | | This study |
| CS152 | *L. monocytogenes* 10403S | $\Delta alsR::alsR$ | | This study |
| JW130 | *L. monocytogenes* 10403S | PrfA* |  | (37) |
| CS445 | *L. monocytogenes* 10403S | $\Delta alsR$ PrfA* | | This study |
| CS456 | *L. monocytogenes* 10403S | $\Delta alsR\Delta alsOP$ | | This study |
| CS175 | *L. monocytogenes* 10403S | $\Delta alsR\Delta lmo0735$ | | This study |
| CS437 | *L. monocytogenes* 10403S | $\Delta alsR\Delta lmo0736$ | | This study |
| CS449 | *L. monocytogenes* 10403S | $\Delta alsR\Delta lmo0737$ | | This study |
| CS166 | *L. monocytogenes* 10403S | $\Delta alsR\Delta lmo0738$ | | This study |
| CS441 | *L. monocytogenes* 10403S | $\Delta alsR\Delta lmo0739$ | | This study |

**Supplemental Table 4.**
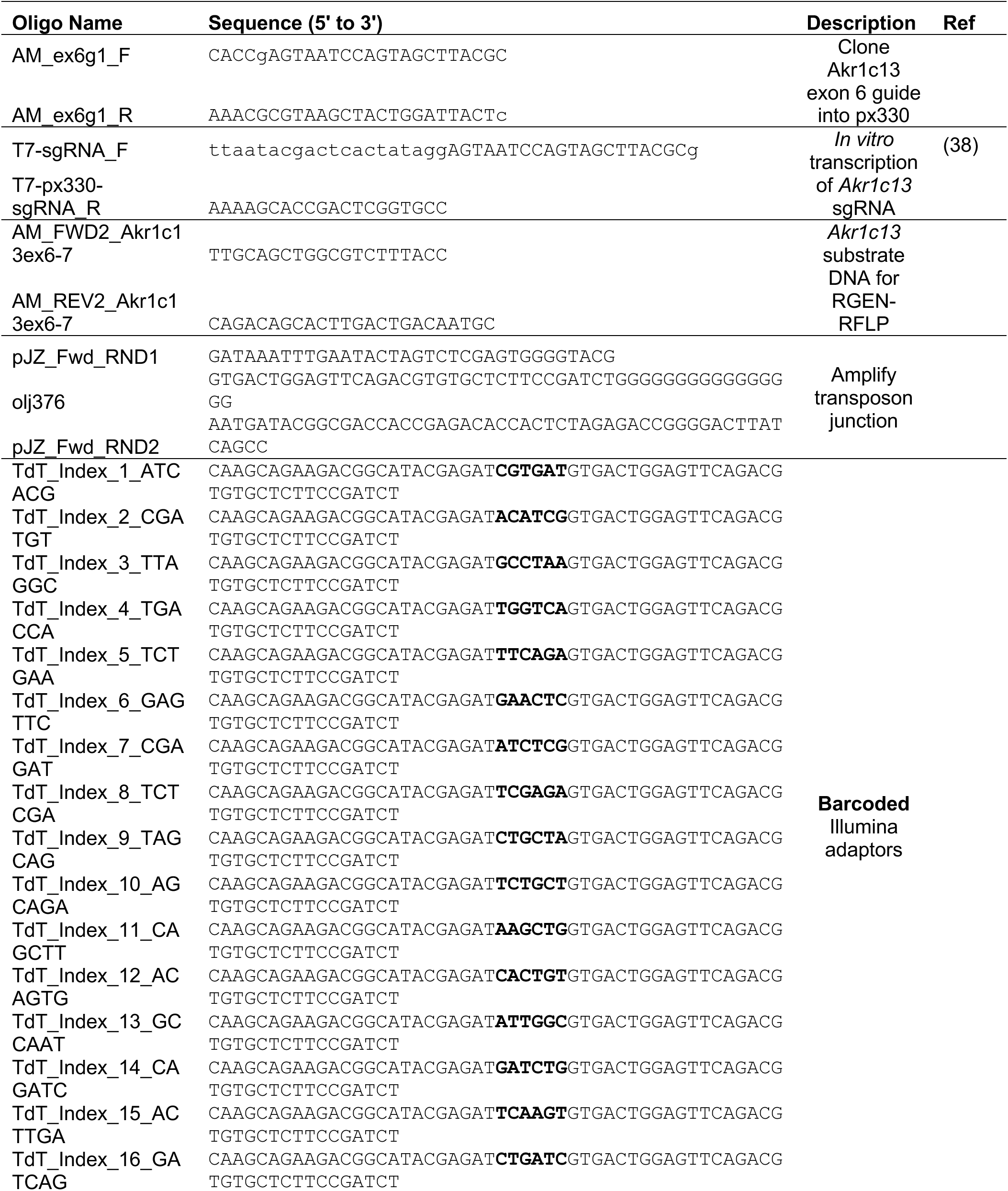
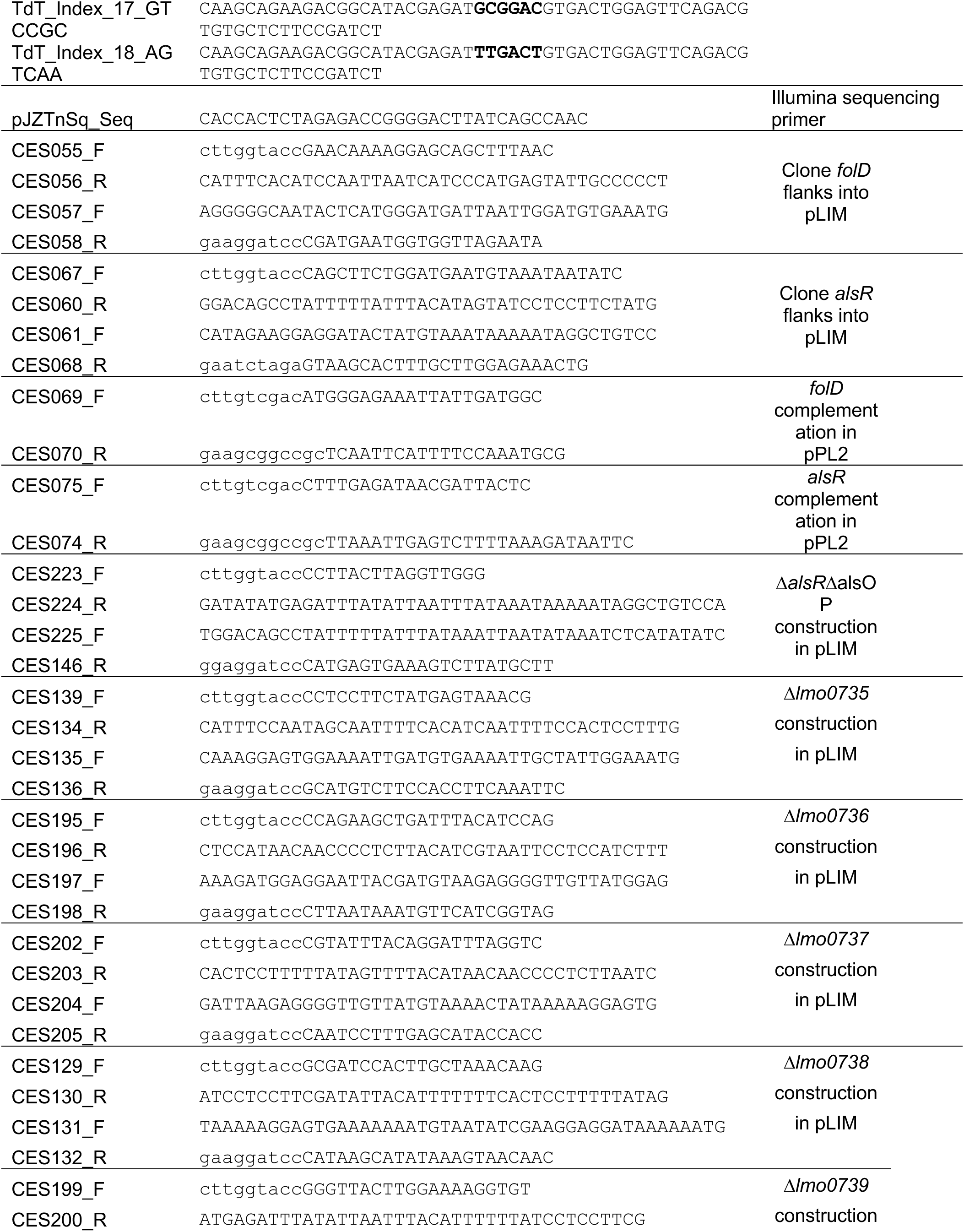
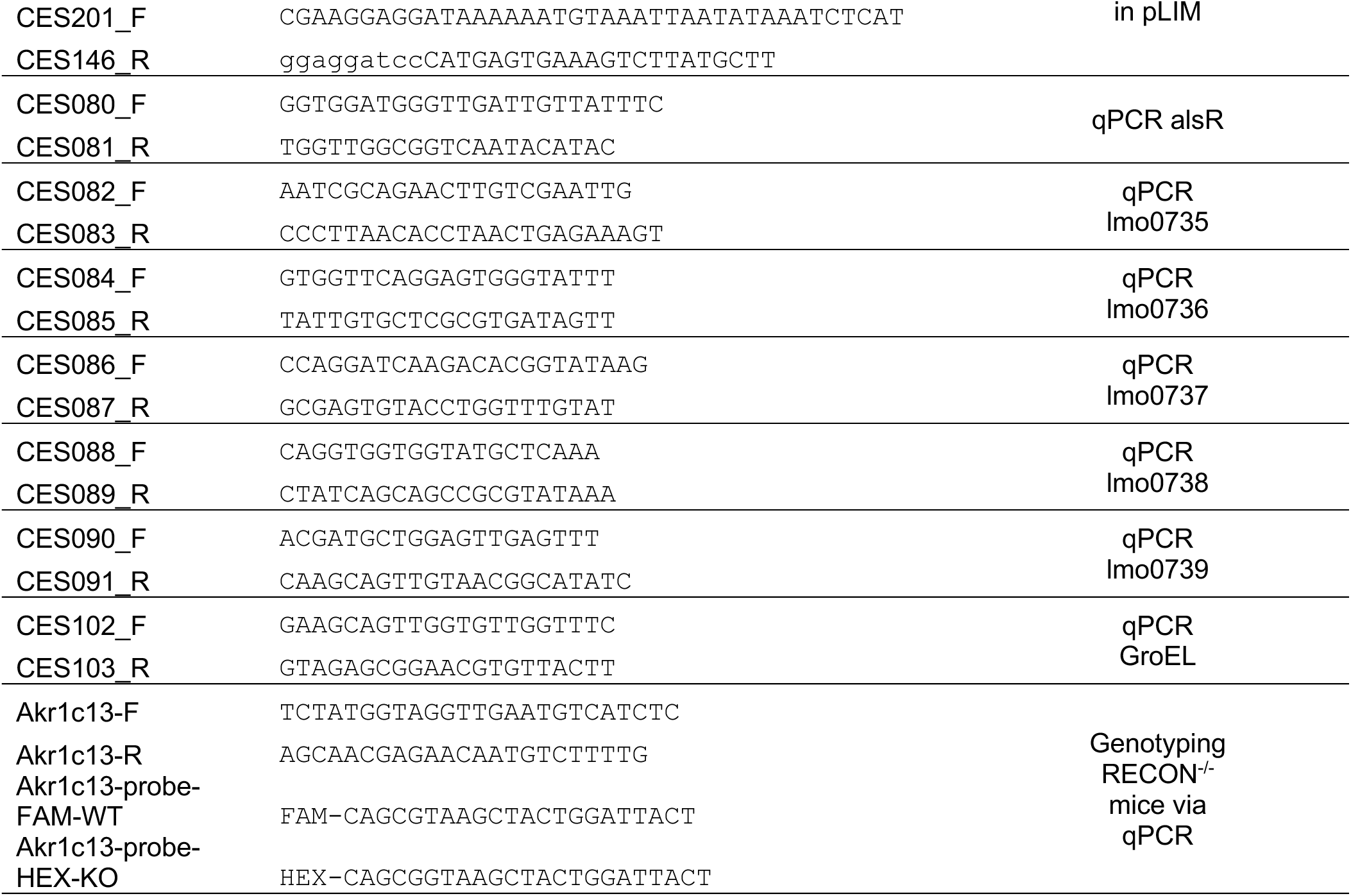
Primers used in this study.

## Supplemental Methods

### SNP PCR genotyping of mice

Mouse genotyping was performed using a custom multiplex *Akr1c13* SNP-based genotyping assay. An end-point PCR was first performed to amplify *Akr1c13* exon 6. The product was diluted 1:1,000 in nuclease-free water, and 1.0 μL was used in a 20 μL qRT-PCR reaction, along with TaqMan Master Mix, primers that amplify *Akr1c13* exon 6 (500 nM final concentration, Table S4) and FAM probes that detect the WT allele mixed with HEX probes that detect the mutant allele (250 nM final probe concentrations, Table S4).

### Bacterial growth curves

Overnight cultures of Lm were back diluted to OD_600_=0.05 and grown at 37°C with shaking until OD_600_ reached 0.4. Bacteria were resuspended in 1X PBS to OD_600_=1.0, diluted 1:100 in BHI or MM and 200 µL of this suspension was distributed to a 96-well plate. The plate was covered in Breathe Easy film (Diversified Biotech #BEM-1) and incubated at 37°C, 237 CPM in a BioTek Synergy HTX plate reader with OD_600_ reads every 30 minutes.

### BMDM growth curves with Lm

BMDM were seeded in 24-well tissue culture treated plates at a density of 2.5 x 10^5^ cells/well with the addition of 100ng/µL murine IFN-γ when indicated and incubated overnight at 37°C 5% CO_2_. Cells were washed once in 1X PBS prior to infection. Overnight cultures of Lm were resuspended in 1X PBS to OD_600_=1.0 and diluted 1:1,000 in DMEM +10% FBS +10% CSF and 500 µL of this infection medium was added to each well (MOI 3). Cells were incubated for 30 minutes to allow phagocytosis, then were washed twice in 1X PBS and replaced with fresh media containing 100µg/mL gentamicin for the remaining time points. At each time point the cells were washed twice in 1X PBS and lysed in 1 mL (0.5, 4, 6 hpi) or 500 µL (2 hpi) ice cold nanopure water. Lysates were diluted in PBS and plated for CFU enumeration on BHI + Streptomycin.

